# Structural insight into the mechanism of 4-aminoquinolines selectivity for the alpha2A-adrenoceptor

**DOI:** 10.1101/561894

**Authors:** Zaibing Li, Jingyu Li, Liyan Liu, Wenyi Deng, Wen Zhang, Huifang Hou, Xinyuan Wang, Zhimei Yang, Xiaoying Wang, Shanze Chen, Yi Wang, Junli Chen, Ning Huang

**Author notes:** Corresponding authors (Junli Chen), (Ning Huang). These authors contributed equally to the work (Zaibing Li, Jingyu Li).

## Abstract

α_2A_-adrenoceptor (AR) is a potential target for the treatment of degenerative diseases of the central nervous system, and α_2A_-AR agonists are one of the most effective drugs for this condition. However, the lack of high selectivity for α_2A_-AR subtype of traditional drugs greatly limits their clinic usage. In this study, a series of homobivalent 4-aminoquinolines conjugated by two 4-aminoquinoline moieties via varying alkane linker length (C2-C12) were characterized for their affinities for each α_2_-AR subtype. Subsequently, docking, molecular dynamics and mutagenesis were applied to uncover the molecular mechanism. Most 4-aminoquinolines (4-aminoquinoline monomer, C2-C6, C8-C10) were selective for the α_2A_-AR over α_2B_- and α_2C_-ARs. Besides, the affinities are of similar linker length-dependence for each α_2_-AR subtype. Among all the compounds tested, C10 has the highest affinity for the α_2A_-AR (p*K*i=-7.45±0.62), which is 12-fold and 60-fold selective over α_2B_-AR and α_2C_-AR, respectively. Docking and molecular dynamics studies suggest that C10 simultaneously interacts with an orthosteric and an “allosteric site” of the α_2A_-AR. The mutation of F205, which is situated at the orthosteric binding pocket decreases the affinity by 2-fold. The potential allosteric residues include Ser90, Asn93, Glu94 and W99. The specificity of C10 for the α_2A_-AR and the potential orthosteric and allosteric binding sites proposed in this study provide valuable guidance for the development of novel α_2A_-AR subtype selective compounds.

## 1. Introduction

alpha2 adrenoceptors (α_2_-ARs) belong to class A rhodopsin-like G-protein coupled receptors, which are sub-classified to α_2A_, α_2B_ and α_2C_-ARs. α_2_-ARs are mainly coupled to Gi protein and inhibit adenylyl cyclase activity, resulting in lower cAMP levels [1].

α_2_-ARs have been gradually recognized as promising antipsychotic therapeutic targets, especially for those associated with affective, psychotic, and cognitive symptoms [2]. α_2A_-AR is widely distributed in central nervous system (CNS), accounting for 90% of α_2_-ARs, and is associated with regulation and strengthen of memory, analgesia, sedation, and has an anti-epileptic effect [3]. Thus, the activation of the α_2A_-AR can improve the clinic symptoms of CNS degenerative diseases. However, increasingly more studies have shown that α_2A_-AR and α_2C_-AR have different or even opposing roles in the CNS [4]. For instance, the activation of the α_2A_-AR ameliorates spatial working memory ability of mice, while the activation of the α_2C_-AR destroys this ability [4]. Thees studies indicate that the selective α_2A_-AR agonists might be one of the most ideal treatments for CNS degenerative diseases. However, the lack of highly selective α_2A_-AR compounds limits the development of α_2A_-AR selective drugs and their clinical use.

In previous studies, homobivalent 4-aminoquinoline compounds (Fig. 1A) were shown to have nanomolar affinity for the α_2_-AR when the three subtypes of the α_2_-AR had not been discovered, and the affinities had tissue-specific differences [5,6]. In addition, the affinities were of linker-length dependence, suggesting there might be a second pocket that is different from the orthosteric pocket (endogenous agonist binding site). Any site on a receptor that is distinct from the orthosteric site is called allosteric site [7]. The development of allosteric modulators has drawn increasingly more attention due to their several potential advantages over traditional (orthosteric) drugs, including having higher subtype selectivity and maintaining spatiotemporal patterns of physiological signaling, etc. [7]. It has been confirmed that α_2_-ARs have three subtypes and have different tissue distributions [1]. The above results suggest that homobivalent 4-aminoquinoline compounds with certain linker lengths might have higher affinity for a certain subtype of α_2_-ARs, which might partly be resulted from the allosteric interaction.

**Figure 1.**
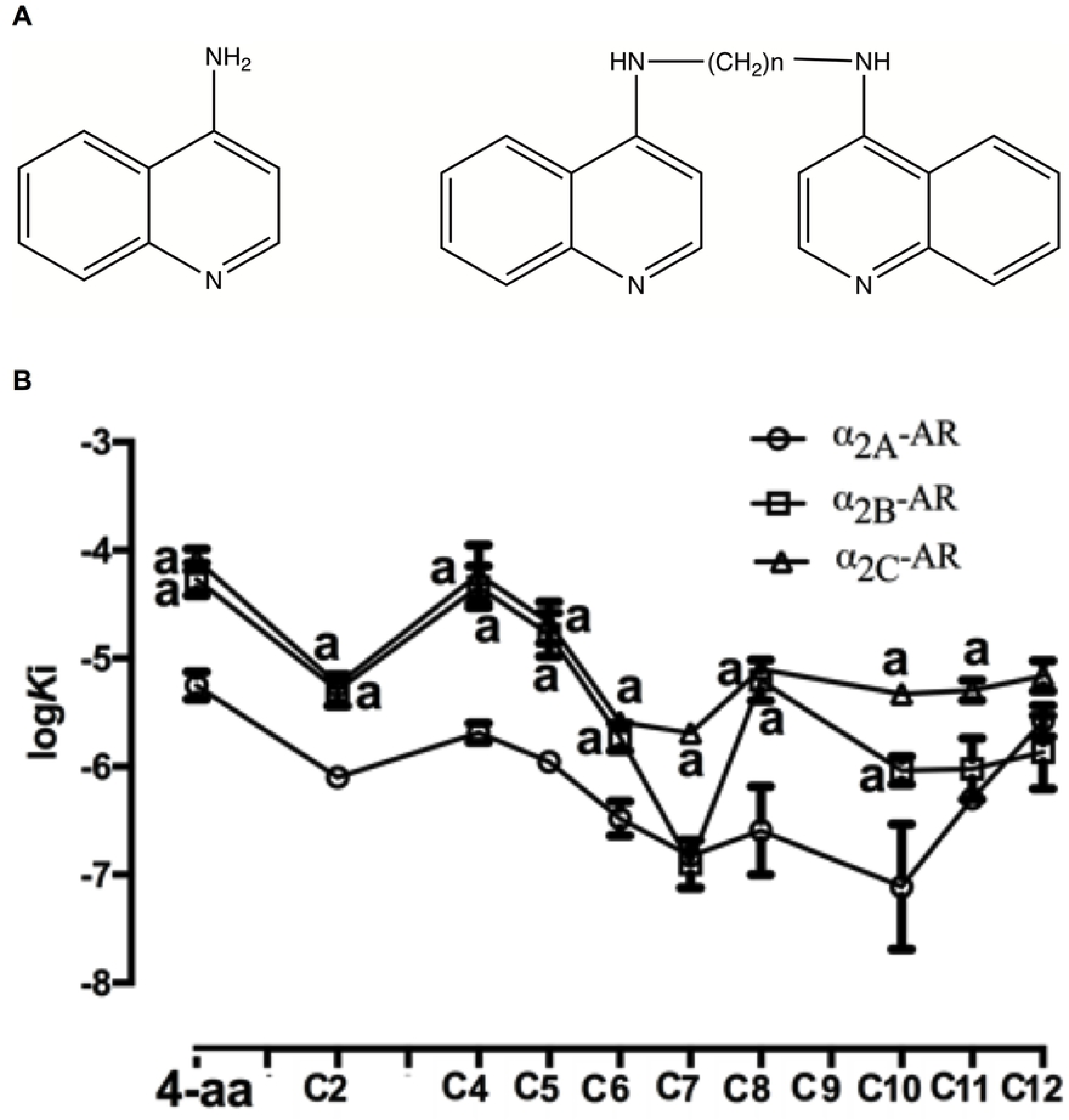
Structures (A) and subtype selective binding (B) of 4-aminoquinoline compounds. Competition binding assay was performed on membranes prepared from α_2A_, α_2B_ or α_2C_-ARs transiently transfected COS-7 cells. All binding curves were fit by a one-site binding model. Affinities were compared using one-way ANOVA and student-Newman-Keuls multiple comparison tests. a: p<0.05 compared to 4-aminoquinoline, b: p<0.05 compared to α_2A-_AR.

In the current study, a series of homobivalent 4-aminoquinoline compounds with different linker length (2-12) were synthesized and their affinities for each α_2_-AR subtype were measured via radioligand binding assays. Molecular docking, molecular dynamics and site-directed mutagenesis studies were then carried out to investigate the interacting sites of 4-aminoquinolines with α_2A_-AR.

## 2. Materials and Methods

### 2.1 Materials

DEAE-Dextran kit was bought from Beyotime Biotechnology (Shanghai, China). [125I] was from PerkinElmer (Waltham, MA, USA). MEM media, Lipofectamine 2000 were from Invitrogen (Carlsbad, CA, USA). IBMX, foskolin, phentolamine, norepinephrine, 1,4-butanediamine, 1,5-diamino-pentane, 1,6-hexanediamine, 1,7-diaminoheptane, 1,8-Oktandiamin, 1,10-Diaminodecane, 1,11-Diaminoundecane, and 1,12-Diaminododecane were from Sigma-Aldrich (St. Louis, MO, USA). Plasmid mini Kit I, Endo-Free Plasmid Maxi Kit were from Omega (Norcross, GA, USA). DMT Enzyme, cAMP-GlO^TM^ Assay kit were from Promega (USA). Human α_2A_-AR, α_2B_-AR and α_2C_-AR in pcDNA3.1^+^ were from Missouri S&T cDNA Resource Center (www.cdna.org).

### 2.2 Synthesis

A series of homobivalent 4-aminoquinoline compounds connected by alkane linkers of varying lengths were synthesized as previously described [8]. All 4-aminoquinolines were dissolved in dimethyl sulphoxide (DMSO). These stock solutions were stored at −80℃.

### 2.3 Cell Culture, transient transfection and membrane preparations

COS-7 cells were cultured in DMEM supplemented with 10% fetal bovine serum and 1% penicillin/streptomycin. Transient transfection of human α_2A_-, α_2B_-, and α_2C_-ARs in pcDNA3.1^+^ vectors was performed using the method described previously [9]. Briefly, 1×10^7^ COS-7 cells were seeded per 150 mm plate and transfected 24 hours later using 14 µg plasmid DNA per plate. 48-72 hours later, cells were scraped into cold PBS, and centrifuged at 500g for 4 minutes at 4°C. The pellet was resuspended in 20 mL of cold solution (20 mM HEPES, 10mM EDTA, pH 7.4). They were disrupted by homogenization for 10 seconds using an ultra-turax homogenizer at 24000 rpm, with an interval of 30 seconds between each homogenization. The homogenate was centrifuged at 1300g for 10 minutes at 4°C and the supernatant was centrifuged at 40,000g for 1 hour at 4°C. The membrane pellet was resuspended in 0.6-1ml buffer (50 mM Tris-HCl, 120 mM sucrose, pH 7.4, and 10% glycerol (v/v)). The membrane suspension was homogenized on ice using an insulin syringe, aliquoted and stored at −80°C. The protein concentration was determined using Bradford reagent.

### 2.4 Site directed mutagenesis

Site-directed mutagenesis was carried out through PCR reaction to mutate the potential orthosteric and allosteric sites of the α_2A_-AR to alanine (A). Primers were shown in supplementary information Table S1. PCR reaction was performed using the gold mix kit, and samples were subjected to 30 cycles of 10 seconds of denaturation at 98°C, 10 seconds of annealing at 55°C, and 7 minutes of elongation at 72°C. A DMT restriction enzyme was used to digest the parental plasmid. The constructed mutant plasmids were transformed into DH5α competent cells. The positive colony with the mutant plasmids was identified by sequencing.

### 2.5. Radioligand binding assays

All ligands and membranes were suspended in buffer containing 50 mM Tris-HCl and 120 mM sucrose. For saturation binding assay, membranes containing each α_2_-AR subtype were incubated with various concentrations of [^125^I]-PIC (0.1-10 nM) in a total volume of 200 µl. For competition binding assay, 4-15 μg of each α_2_-AR subtype membranes were incubated with 400 pM of [^125^I]-PIC and increasing concentrations of test compounds in a total volume of 200 mL. Non-specific binding was defined as binding in the presence of 100 µM phentolamine. All the reaction mixtures for all binding experiments were incubated at room temperature for 1h, and the reaction was terminated by PBS and vacuum filtration through GF/B filters. Radioactivity was measured by a [^125^I] beta counter.

### 2.6 cAMP assay

The production of intracellular cAMP was determined using cAMP-GlO^TM^ Assay kit. Briefly,1×10^5^ cells ml^-1^ of transiently transfected COS-7 cells were seeded into 96 well plates and cultured for 48 h. Cell were then washed with DMEM and were treated for 30 minutes with 20 µL DMEM containing 1 mM IBMX, 30 µM forskolin and different concentrations (10^-10^-10^-3^ M) of test compounds. Cells were lysed for 30 minutes at room temperature in cAMP-Glo lysis buffer (20 µL per well). 40 µL/well detection solution (containing PKA substrate and PKA holoenzyme) was added and cultured at room temperature for 20 min. 80 µL/well of Kinase-Glo^TM^ Regeat was added, which were cultured for 10 minutes at room temperature. Luminescence was measured with luminometers.

### 2.7 Homology modeling

The human α_2A_-AR sequence obtained from the uniport database was used to screen the experimentally modeled protein structure and the human β1-AR (PDB ID: 2YCY) crystal structure was selected as the template. The homology model of the α_2A_-AR was built using the sequence alignment of α_2A_-AR and β1-AR via SWISS-MODEL server. The predicted 3D structure of the α_2A_-AR was further optimized by MODELLER (v9.16), which was then evaluated using ERRAT plot and Ramachandran plot.

### 2.8 Molecular Docking

Molecular docking was performed using Autodock vina to investigate the binding mode between 4-aminoquinolines and the α_2A_-AR. The search grid of the α_2A_-AR was identified as center_x: 18.16, center_y: 20.7, center_z: −6.67 with dimensions size_x: 100, size_y: 100, and size_z:100. All other parameters were set as “default”. The best-scoring pose as judged by the Vina docking score was chosen and visually analyzed using Accelrys Discovery Studio Client version 3.1 (Accelrys Software Inc. USA).

### 2.9 Molecular dynamics (MD) simulations

The MD simulations were performed in a hydrated dipalmitoyl phosphatidylcholine (DPPC) bilayer using the GROMACS (version 2016) software package [10]. CHARMM36 force field [11] was used for protein and lipid, and CGenFF force field [12] was used for small molecules. The system was heated to 300k for 5 ns in NVT ensemble. Subsequently, the equilibration simulation ran for 10 ns in NPT ensemble (Pressure=1atm and temperature=300k). Finally, the production simulation was conducted for 50 ns. RMSD and C-αRMSF analyses were performed to monitor the stability of the system, and the binding free energy was calculated via MM/PBSA software.

### 2.10 Data analysis

Nonlinear regression analysis of saturation, competition binding, and inositol phosphate accumulation assay data was performed using GraphPad Prism 6.0 (San Diego, CA, USA). Statistically significant differences (p<0.05) between the affinities of all compounds were compared using one-way ANOVA and student-Newman-Keuls multiple comparison tests.

## 3. Results and discussion

### 3.1 The affinity of 4-aminoquinolines at α_2_-ARs

A series of 4-aminoquinolines with different linker lengths (2-12) shown in Fig.1A were synthesized as described previously [8]. All synthesized compounds were identified by proton nuclear magnetic resonance and mass spectrometry (Supplementary Information, Fig. S1&S2).

To address subtype selectivity of 4-aminoquinolines, their apparent affinities were evaluated on membrane-expressed human α_2_-ARs via competition radioligand binding assays using [^125^I]-PIC (α_2A_-AR *K*_D_= 0.2090±0.06 nM, α_2B_-AR *K*_D_= 0.6020±0.081 nM, α_2C_-AR *K*_D_= 0.8023±0.053 nM).

In general, 4-aminoquinoline compounds showed nano- to sub-micromolar affinities for the three α_2_-AR subtypes (Fig. 1B, Table 1). Most 4-aminoquinoline compounds had greater affinity than the 4-aminoquinoline monomer for each α_2_-AR subtype (Fig. 1B, Table 1), suggesting that conjugation of two 4-aminoquinoline pharmacophores can increase the affinity for the α_2_-ARs. This is consistent with previous studies [13]. The 4-aminoquinoline and homobivalent 4-aminoquinolines with linker lengths between 2-6, and 8-10 carbons showed a 6-24 fold selectivity for α_2A_-AR over α_2B_-AR (p<0.05); and all the 4-aminoquinoline compounds, were of 2.6-60 fold selectivity for α_2A_-AR over α_2C_-AR (p<0.05), suggesting that the molecular size of these 4-aminoquinoline compounds can fit better for the ligand binding site of the α_2A_-AR.

**Table 1.** Binding affinities of homobivalent 4-aminoquinolines for α_2_-ARs. All data presented are the mean±SE of separate assays, performed in duplicate. *K*i refers to the concentration of ligand required to occupy 50% of unoccupied receptors, calculated according to the Cheng-Prusoff equation: *K*i=IC_50_/1+([L]/*K*_D_) where [L] is the radioligand concentration and *K*_D_ is the dissociation constant. p*K*i is the negative log of the *K*i value. ^a^ p<0.05 compared to 4-aminoquinoline.

| Compound | $\alpha_{2A}$-AR | | | $\alpha_{2B}$-AR | | | $\alpha_{2C}$-AR | | |
| --- | --- | --- | --- | --- | --- | --- | --- | --- | --- |
|  | p*K*<sub>i</sub> | *K*<sub>i</sub>(nM) | n | p*K*<sub>i</sub> | *K*<sub>i</sub>(nM) | n | p*K*<sub>i</sub> | *K*<sub>i</sub>(nM) | n |
| 4-aminoquinoline | -5.25±0.12 | 5596 | 3 | -4.27±0.15<sup>b</sup> | 53790 | 3 | -4.09±0.10<sup>b</sup> | 81470 | 3 |
| C2 | -6.10±0.07 | 797 | 3 | -5.30±0.14<sup>b</sup> | 5075 | 3 | -5.24±0.06<sup>b</sup> | 5750 | 3 |
| C4 | -5.69±0.09 | 2062 | 3 | -4.34±0.19<sup>b</sup> | 45870 | 3 | -4.22±0.27<sup>b</sup> | 60440 | 3 |
| C5 | -5.96±0.06 | 1109 | 3 | -4.78±0.20<sup>b</sup> | 16560 | 3 | -4.67±0.19<sup>b</sup> | 21320 | 3 |
| C6 | -6.52±0.18 | 329.4 | 3 | -5.73±0.13<sup>b</sup> | 1883 | 3 | -5.59±0.04<sup>b</sup> | 2585 | 3 |
| C7 | -6.83±0.03 | 149 | 3 | -6.90±0.22 | 126 | 3 | -6.69±0.04<sup>b</sup> | 2042 | 3 |
| C8 | -6.78±0.44 | 259 | 3 | -5.20±0.19<sup>b</sup> | 6266 | 3 | -5.10±0.08<sup>b</sup> | 7898 | 3 |
| C10 | -7.45±0.62 | 78 | 3 | -6.04±0.13<sup>b</sup> | 923 | 3 | -5.33±0.01<sup>b</sup> | 4697 | 3 |
| C11 | -6.30±0.01 | 506 | 3 | -6.02±0.28 | 951 | 3 | -5.29±0.09<sup>b</sup> | 5080 | 3 |
| C12 | -5.58±0.15 | 2640 | 3 | -5.87±0.34 | 1354 | 3 | -5.16±0.14 | 6857 | 3 |

There is a similar linker length-affinity relationship for each α_2_-AR subtype. At the α_2A_-AR, there are two domains of high affinity for 4-aminoquinolines, that is when the linkage comprises 2 and 10 methylene groups (Fig. 1B, Table 1). Specifically, C2 had an affinity of 797 nM, which decreased when the linker length was increased to 4 or 5 (Fig. 1B, Table 1). We speculated that C2 has the most suitable size to fit the binding pocket of the α_2A_-AR, and therefore forms more interactions with the receptor. In contrast, the size of 4-aminoquinolie is too small, while that of C4 and C5 is too big which might lead to steric clashes, resulting in decreased affinity. Interestingly, the affinity increased afterwards when the linker of homobivalent 4-aminoquinoline compounds was further lengthened, reaching at the peak at 10 (78 nM) carbon atoms, which decreased thereafter when the linker was further lengthened to 11 and 12 (Fig. 1B, Table 1). We hypothesized that the linker length of 6-10 is of appropriate length for the compounds to interact with the residues that are located at the extracellular surface of the receptor. The homobivalent 4-aminoquinoline compounds had a similar trend of linker length-affinity relationship for α_2B_-AR and α_2C_-AR, but the peak was located at C2 (α_2B_-AR, 5075; α_2C_-AR 5750 nM) and C7 (α_2B_-AR, 125 nM; α_2C_-AR, 2042 nM) (Fig. 1B, Table 1).

### 3.2 The effect of C10 on the production of cAMP

α_2_-ARs are mainly coupled to Gαi protein and inhibit adenylyl cyclase to produce cAMP. However, α_2_-ARs could also increase cAMP accumulation by either activating adenylyl cyclase II or under lower agonist concentrations [14, 15].

To define the effect of C10 on cAMP production in COS-7 cells, cAMP accumulation assay was carried out by testing the effect of C10 on foskolin induced activation of α_2A_-AR. C10 showed a biphasic effect on cAMP production in COS-7 cells expressing the α_2A_-AR. At lower concentrations (10^-10^-10^-7^ M), C10 increased cAMP production, whereas at higher concentrations (10^-6^-10^-4^ M) inhibited cAMP accumulation (Fig. 2A). This biphasic effect is similar to previously reported results with epinephrine in CHO cells, which showed an inhibition effect from 1 to 10 nM, and an increase from 100 to 1000 nM [15]. In COS-7 cells, we have also shown that norepinephrine inhibited the production of cAMP at lower concentrations (10^-10^-10^-8^M), while stimulated the expression of cAMP at higher concentrations (10^-3^-10^-7^M) (Fig. 2B). As reported previously [15], phentolamine, a known antagonist of the α_2A_-AR, did not have any effect on the production of cAMP (Fig. 2C), but suppressed NA to produce cAMP, with an IC50 value of 97.05±0.086 nM (Fig. 2D).

**Figure 2.**
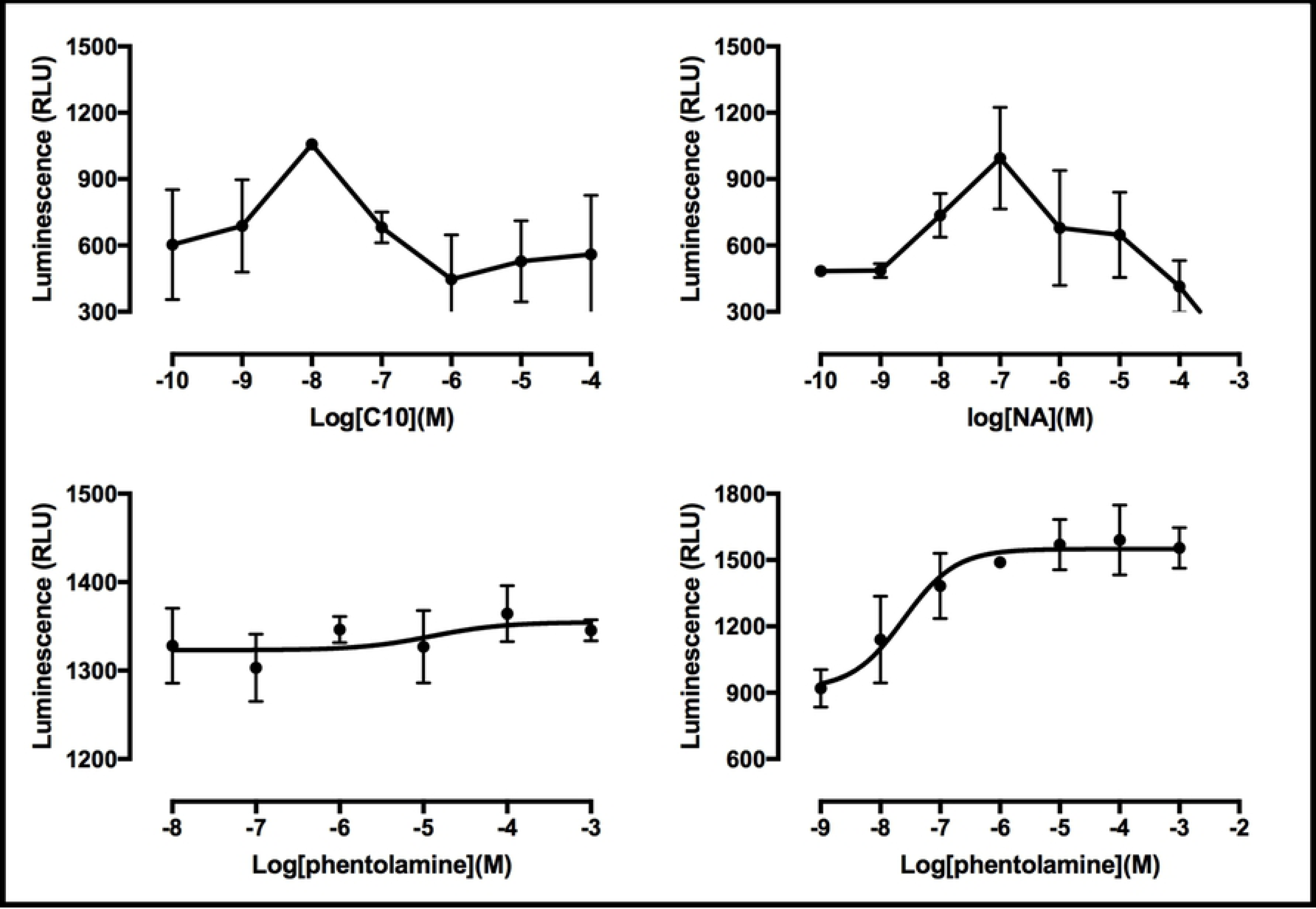
The effect of C10 on foskolin induced cAMP accumulation. COS-7 cells transiently transfected with α_2A_-AR were treated for 30 min at 37℃with 20 µL DMEM containing 1 mM IBMX, 30 µM forskolin and increasing concentrations of C10 (A), norepinephrine (B), phentolamine (C), phentolamine and 10^-4^ M norepinephrine (D). cAMP concentration was monitored with luminometers.

### 3.3 The interactions predicted by molecular docking

Given that 4-aminoquinoline monomer and C10 showed specificity for α_2A_-AR, and C10 had the highest affinity for the α_2A_-AR among all 4-aminoquinolines, their possible interacting sites were predicted via molecular docking. The α_2A_-AR homology model was firstly constructed based on the β1-AR (PDB ID: 2YCY) template via swiss-model. The predicted 3D structure was further optimized by MODELLER (v9.16), which was then evaluated using ERRAT plot and Ramachandran plot. The ERRAT program gave a score of 90.862, and the results of the Ramachandran plot showed that 97% residues were in the most favored and allowed regions, indicating a good quality model.

Molecular docking results showed that 4-aminoquinoline monomer formed a hydrogen bond with Asp113 (TMIII) (Fig. 3A), which has been shown to be a conserved orthosteric residue (endogenous agonist binding site) among aminergic GPCRs [16]. The results suggest that 4-aminoquinoline monomer is situated at the orthosteric binding pocket of the α_2A_-AR. Interestingly, one quinoline moiety of C10 is within the orthosteric binding pocket, forming a π-π interaction with Trp387, while the other quinoline ring is located at the extracellular surface of the receptor, forming a π-π interaction with Trp99 (ECL1) and a hydrogen bond with Cys106 (TMIII) (Fig. 3B).

**Figure 3.**
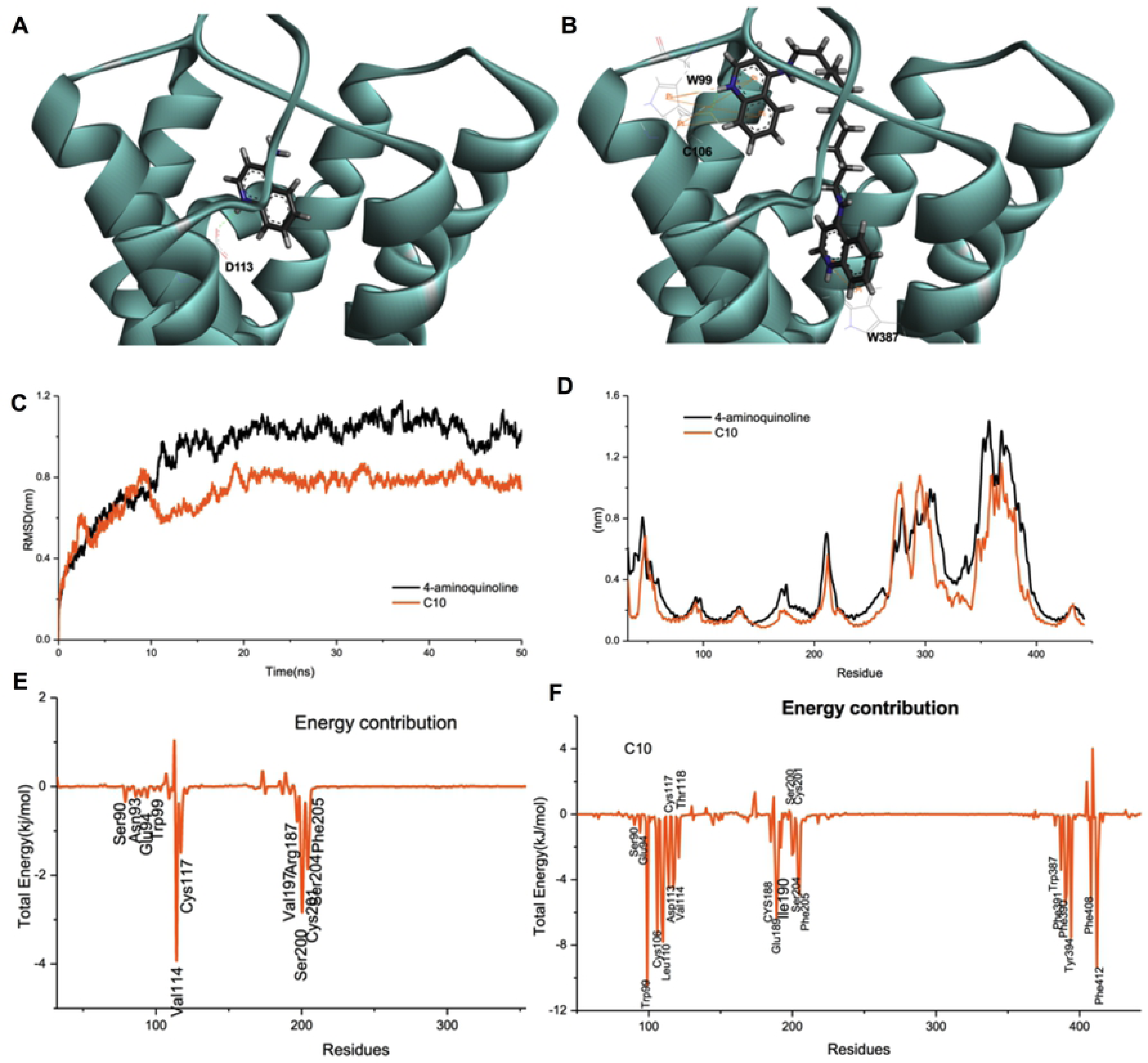
The interactings between 4-aminoquinoles and the α_2A_-AR. 4-aminoquinoline monomer (A) or C10 (B) was docked into the α_2A_-AR homology based on a human β1-AR crystal structure (2YCY). Molecular dynamics was subsequently performed. The root-mean-square deviation (RMSD) was shown in C, and the root mean square fluctuation (RMSF) profiles were in D. And each residue energy contribution to the binding free energy of the system was in E (4-aminoquinoline-α_2A_-AR) and F (C10-α_2A_-AR).

### 3.4 The interactions predicted by molecular dynamics simulations

A 50 ns molecular dynamics simulation was carried out to further validate the reliability of the docking results between 4-aminoquinolines and the α_2A_-AR. The root-mean-square deviation (RMSD) of each system showed that the α_2A_-AR docked with either 4-aminoquinoline monomer or C10 achieved equilibrium at around 20 ns with RMSD average value of approximately 0.8 nm and 1 nm, respectively (Fig. 3C). The root mean square fluctuation (RMSF) profiles were then analyzed to investigate the fluctuations of residues with the conformational transition. We found that the 4-aminoquinoline monomer-α_2A_ and C10-α_2A_ complexes had very similar RMSF profiles. The RMSF was around 1.1 nm at the extracellular and intracellular loops, and was less than 0.5 nm for other residues, suggesting the loops are relatively more fluctuant than the alpha helices and beta sheets (Fig. 3D).

The binding free energy of the protein-compound complexes was calculated using MM/PBSA method. 4-aminoquinoline monomer demonstrated negative binding free energy value of −111.69 KJ/mol, while C10 possessed value of −289.70 KJ/mol, indicating C10 has higher affinity for the α_2A_-AR than the 4-aminoquinoline monomer. This is in a good agreement with the *K*i values of our competitive binding assays, which supports the reliability of our force field parameters and MD simulations.

g_mmpbsa was applied to determine each residue contribution to the binding free energy to determine the key residues interacting with the 4-aminoquinolines. Fig. 3E and Table S2 demonstrated that the key residues of the 4-aminoquinoline-α_2A_ complex that contributed to the total free energies were located at TMⅢ (Val114， Cys117), TMⅤ (Val197, Ser200, Cys201, Ser204, Phe205), TMⅥ (Trp387, Phe390, Phe391， Tyr394), TMⅦ (Phe412), ECL2 (Arg187). Interestingly, some residues which are located at the extracellular part of TM II or ECL1, also contributed to the binding free energy but with lower energies, including Ser90 (TMⅡ, -0.27 KJ/mol), Asn93 (TM Ⅱ, −0.06 KJ/mol), Glu94 (TM II, −0.35 KJ/mol) and Trp99(ECL1, −0.13 KJ/mol) (Fig. 3E, Table S2).

C10-α_2A_ complex had very similar features to 4-aminoquinoline-α_2A_ complex but with higher energy contributions of each residue. The key residues of the C10-α_2A_ complex that contributed more than 2 KJ/mol were located at TMⅢ (Cys106, Tyr109, Leu110, Asp113, Val114, Cys117, Thr118,)， TMⅤ (Ser200, Cys201, Ser204, Phe205), TMⅥ (Trp387, Phe390, Phe391, Tyr394), TMⅦ (Phe408, Phe412, Phe413), ECL2 (Cys188, Glu189, Ile190) (Fig 3F, Table S2). Similar to 4-aminoquinoline-α_2A_ complex, some residues in the C10-α_2A_ complex at the extracellular part of TMⅡor ECL1 also contributed to the binding free energy with lower energies, including Glu94 (TMⅡ, −1.10 KJ/mol), Ser90 (TMⅡ, −0.64 KJ/mol), Asn93 (TMⅡ, −0.53 KJ/mol) and Trp99 (ECL1, −10.48 KJ/mol) (Fig. 3F, Table S2).

The endogenous agonist binding pocket of aminergic GPCRs is called the orthosteric site. It has been shown previously that Ser200, Cys201, Ser204 of the α_2A_-AR interact with catecholamines, and are involved in the activation of the receptor [17], suggesting Ser200, Cys201 and Ser204 are located at the orthosteric binding site of the α_2A_-AR. The interaction of 4-aminoquinoline and C10 with these three resides indicate that 4-aminoquinoline and C10 are within the orthosteric binding pocket of the α_2A_-AR. Besides, our results show that the key residues that contribute to the total free energies are located at the transmembrane regions of helixes Ⅲ， Ⅴ， Ⅵ， Ⅶ, and ECL2， which are consistent with the reported orthosteric binding pocket of aminergic GPCRs [18,19].

The development of allosteric modulators has been shown to be an effective way to obtain subtype selective ligands [7]. Previous studies have suggested that there is a “common” allosteric site among different aminergic GPCRs, which comprises the residues from ECL2 and the extracellular part of TMII and TMVII [20, 21]. In the present study, molecular dynamics simulations of both 4-aminoquinoline-α_2A_ and C10-α_2A_ complexes demonstrate that some residues which are located at ECLI or the extracellular part of TMII also contributed to the total free energies, including Trp99 (ECLI), Ser90, Asn93 and Glu94 (TMII). These results suggest that Trp99, Ser90, Asn93 and Glu94 might be allosteric sites of the α_2A_-AR. The lower energies contributed by the potential allosteric sites compared to the orthosteric residues could be due to the fact that allosteric modulators normally have lower affinities than the orthosteric ligands [22]. The interaction of one 4-aminoquinoline moiety with the allosteric site might cause the structural changes of the α_2A_-AR, thus leading to the higher affinity and selectivity of C10 for the α_2A_-AR.

### 3.5 F205A decreased the affinity of C10 for the α_2A_-AR

In order to investigate the molecular mechanism of the specificity of C10 for the α_2A_-AR, two potential orthosteric residues (Glu189, Phe205) and allosteric sites (Asn93, Trp99) predicted by molecular dynamics simulations, were selected and mutated to alanine. Subsequently competition binding assay was performed to investigate the effect of the mutants on the affinity of C10 for the α_2A_-AR. We found that F205A reduced the affinity by 2-fold (Fig. 4A, Fig. 4B, Table 2), indicating F205 plays an important role during the interaction of C10 and the α_2A_-AR.

**Figure 4.**
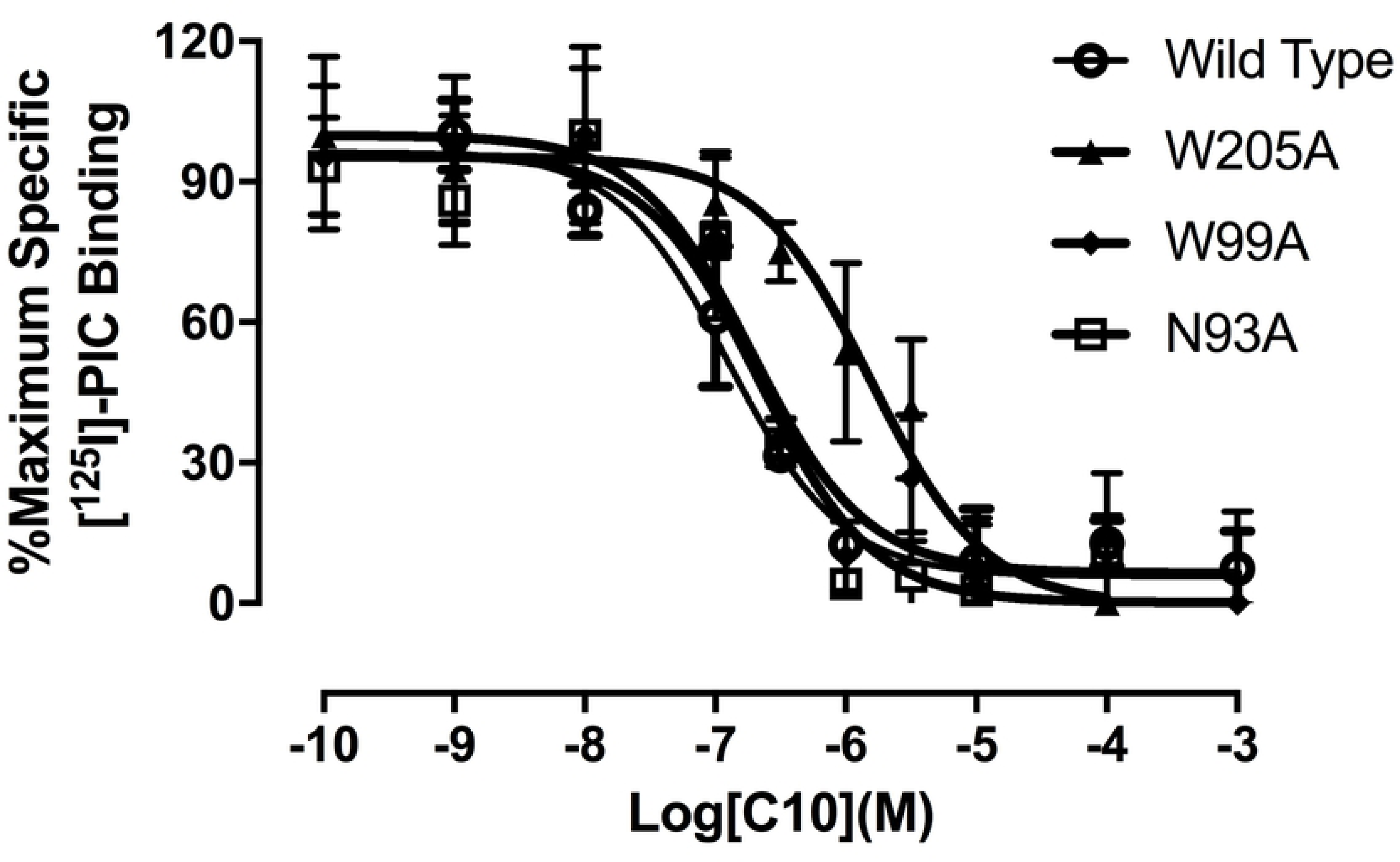
Competitive binding affinity of C10 for α_2A_-AR mutants. Competition by C10 for specific binding of 400pM [^125^I]-PIC (400 pM) to membranes prepared from wild type α_2A_-AR, or α_2A_-AR mutants, F205A, W99A, and N93A transfected COS-7 cells. Points are mean percentage of maximum specific binding and vertical bars represent standard error. Curves were best fit to a single-site model. Affinities were compared using one-way ANOVA and student-Newman-Keuls multiple comparison tests.

**Table 2.**
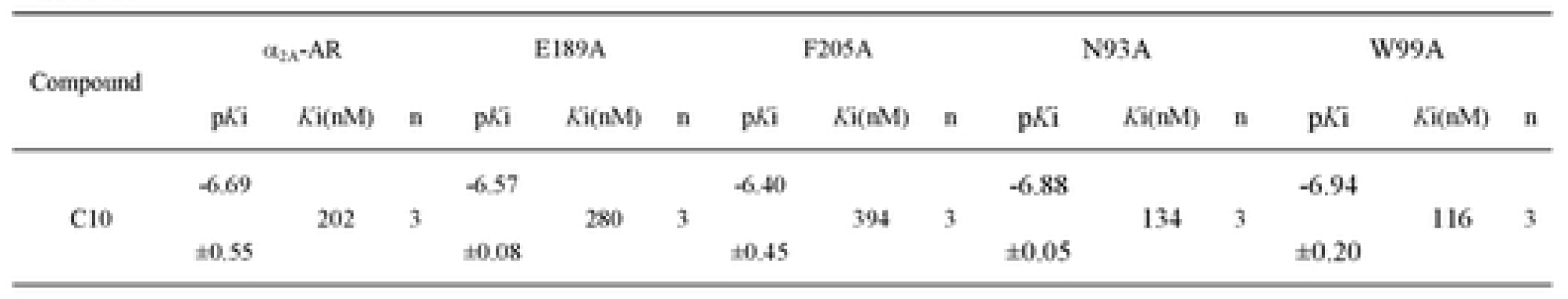
Binding affinities of C10 for α_2A_-AR mutants. All data presented are the mean±SE of separate assays, performed in duplicate. *K*i refers to the concentration of ligand required to occupy 50% of unoccupied receptors, calculated according to the Cheng-Prusoff equation: *K*i=IC_50_/1+([L]/*K*_D_) where [L] is the radioligand concentration and *K*_D_ is the dissociation constant. p*K*i is the negative log of the *K*i value.

Asn93 and Trp99 are potential allosteric sites of the α_2A_-AR predicted by our docking and dynamics studies. However, both W99A and N93A did not decrease the affinity for C10. (Fig. 4, Table 2). This could be because that allosteric modulators normally have lower affinities at the allosteric site than the orthosteric ligands for the orthosteric pocket as discussed above [22, 23]. Accordingly, the mutation of one residue of the allosteric pocket may not produce a significant effect on the affinity of C10 for the whole receptor. Thus, double mutations and association kinetics or dissociation kinetics assays are required to further validate whether Asn93 and Trp99 are truly the binding site of C10.

In this study, we demonstrated that homobivalent 4-aminoquinolines have higher affinity for the α_2A_-AR over α_2B_-, α_2C_-ARs. Specifically, C10 has the highest affinity for the α_2A_-AR among all the 4-aminoquinolines and Phe205 has been confirmed to be one of the interacting sites. Most importantly, we have for the first time proposed the potential allosteric site of the α_2A_-AR. This study will provide valuable structural information for the development of novel α_2A_-AR subtype selective compounds.

## Acknowledgements

This work was supported by grants from the National Natural Science Foundation of China (81803648, 81470931, 31401188, 31701098, 81803183) and Sichuan University (2018SCUH0039, 2016SCU11021, 2016SCU04A08).

